# Characterization of the *Rosa roxbunghii* Tratt transcriptome and analysis of *MYB* genes

**DOI:** 10.1101/392720

**Authors:** Xiaolong Huang, Huiqing Yan, Lisheng Zhai, Zhengting Yang, Yin Yi

**Affiliations:** Key Laboratory of Plant Physiology and Development Regulation, school of life science, Guizhou Normal University, Guiyang, China; Karst Key Laboratory of biodiversity conservation in Southwest China, National Forestry Administration, Guiyang, China

**Keywords:** *Rosa roxbunghii* Tratt, transcriptome, MYB transcription factor, EST-SSR

## Abstract

*Rosa roxbunghii* Tratt belongs to the Rosaceae family, and the fruit is flavorful, economic, and highly nutritious, providing health benefits. MYB proteins play key roles in *R. roxbunghii*’ fruit development and quality. However, the available genomic and transcriptomic information are extremely deficient. Here, a normalized cDNA library was constructed using five tissues, stem, leaf, flower, young fruit, and mature fruit, with three repetitions, and sequenced using the Illumina HiSeq 2500 platform. *De novo* assembly was performed, and 470.66 million clean reads were obtained. In total, 63,727 unigenes, with an average GC content of 42.08%, were determined and 59,358 were annotated. In addition, 9,354 unigenes were assigned the Gene Ontology category, and 20,202 unigenes were assigned to 25 Eukaryotic Ortholog Groups. Additionally, 19,507 unigenes were classified into 140 pathways of the Kyoto Encyclopedia of Genes and Genomes database. Using the transcriptome, 18 candidate *MYB* genes that were significantly expressed in mature fruit, compared with other tissues, were obtained. Among them, 10 R2R3 *MYB* and 1 R1 *MYB* were identified. The expression levels of 12 *MYB* genes randomly selected for qRT-PCR analysis were consistent with the RNA-seq results. A total of 37,545 microsatellites were detected, with an average EST-–SSR frequency of 0.59 (37,545/63,727). This transcriptome data will be valuable for identifying genes of interest and studying their expression and evolution.

## Introduction

*Rose roxburghii* Tratt, a fruit crop and deciduous horticultural shrub, belongs to the Rosaceae family and is mainly distributed in southwestern China. It has several nutritional and functional components, including polysaccharides, flavonoids, troterpenes, and superoxide dismutase activity [1, 2]. Current pharmacological research indicates that it could be used for treatments of arosclerosis and senescence-retardation. In addition, its fruits have radio protective, anti-tumor, antimutagenic, and genoprotective activities [3–5]. As a traditional Chinese medicine, it can be further processed into fruit juice, preserves, and fruit wine [6]. MYB proteins play key roles in fruit development and quality [7, 8]. However, genes of the *R. roxburghii MYB* superfamily have not been comprehensively identified and evaluated. Consequently, we located and characterized *R. roxburghii MYB* genes.

Among the different transcription factor families, MYBs form the largest and most functionally diverse superfamily, and they are involved in regulating cell activities and plant development. The N-terminus of a MYB domain is composed of adjacent tandem repeats [8]. The repeat encodes 50–53 amino acid residues and contains helices forming a helix-turn-helix domain that interact with the major grooves of specific DNA sequences [9]. MYB superfamily members can be classified into several subfamilies, including R1 MYB, R2R3 MYB, R1R2R3 MYB, 4R MYB, and atypical MYB-like proteins [10]. Many MYB superfamily proteins and their functions have been determined in different species. Due to the conserved domains, the R2R3 MYB type is predominant in plants [11]. MYBs are involved in regulating plant growth, development, and stress resistance, including the anthocyanin biosynthetic pathway, trichome initiation and development, flavonoid or phenylpropanoid metabolism, secondary wall biosynthesis, sugar signaling and responses to abiotic or biotic stress [12, 13].

Genomic information is not presently available for *R. roxburghii* in an online database. Thus, high-throughput Illumina sequencing could be performed to determine new transcripts, gene expression levels, and an accurate transcriptome profile [14]. Now it becomes a powerful methodology for species that lack reference genome information. Assembled unigenes with different database annotations can be assayed to evaluate genetic characteristics and metabolic pathways. Recently, fruits of *R roxburghii* at three different developmental stages were collected and analyzed using Illumina sequencing technology. A previous study generated 106,590 unigenes using *de novo* assembly. Using a BLAST algorithm-based search, 9,301 and 2,393 unigenes were categorized into Gene Ontology (GO) and Clusters of Orthologous Group (COG) categories, respectively. Additionally, 7,480 unigenes were assigned to 124 pathways in the Kyoto Encyclopedia of Gene and Genome (KEGG) pathway database. Among the unigenes, 9,131 potential EST–SSR (simple sequence repeat) loci were identified [15].

To better understand the profiles of different tissues in *R. roxburghii,* leaf, stem, flower, young fruit and mature fruit were collected. In this study, the Illumina platform was used to construct a cDNA library using 15 mixed tissues to obtain transcriptome information. MYBs, which are significantly expressed in mature fruit, were identified. For verification, randomly selected *MYB* genes were analyzed by qRT-PCR. EST–SSRs were subjected to a genetic diversity analysis of *R. roxburghii.* The obtained information provides a valuble resource for functional gene analyses, in particular of *MYB* genes.

## Materials and methods

### Biological materials

Wild *Rose roxburghii* were planted in Guizhou Normal University, Guizhou, China. Different tissues were collected including leaf, stem, flower, young fruit (50 days after flowering, YF) and mature fruit (120 days after flowering, MF). Each tissue was in three independent experiments. These materials were immediately frozen in liquid nitrogen, then mechanically grounded into fine powder and stored at −80 °C for further experiment.

### RNA extraction and sequencing

A total of 1 g of frozen tissues from leaf, stem, flower, young fruit and mature fruit were weighed. RNA was isolated using the Trizol method (Takara, Japan), flowing the manufacturer’s guidelines. RNA quality was determined with Nanodrop (Wilmington, USA) and Agilent 2100 (Santa Clara, USA). Fifteen libraries were created; three for each of the individual tissues or growth stage. Samples of mRNA were selected and randomly fragmented. Using these fragments as templates, cDNA was transformed and purified. The adapters were connected to distinguish different sequencing samples. Strand specific cDNA libraries were constructed for sequencing through Illumina HiSeq 2500 (Illumina, San Diego, USA).

### De novo assembly and functional annotation

The clean reads were selected from raw data with filtering out adaptor-only reads, reads with more than 5% N bases unknown, and low-quality reads (reads containing more than 50% bases with Q-value≤10). Trinity assembly program was used to obtain data. To determine the function of the unigenes, BLASTx alignment with an E-value cut off of 10^-5^ was performed with different database, including KOG (Eukaryotic Ortholog Groups, http://www.ncbi.nlm.nih.gov/KOG/) [16], Nr (NCBI non-redundant protein database, http://www.ncbi.nlm.nih.gov/), KEGG (Kyoto Encyclopedia of Genes and Genomes, http://www.genome.jp/kegg/) [17], Gene Ontology (GO, http://www.geneontology.org/) and Swiss-Prot (http://www.expasy.ch/sprot) [18, 19]. The best-aligning results were used to identify sequence direction of the unigenes. If different databases conflicted, the results were prioritized in the following order: Nr, Swiss-Prot, KEGG, COG and GO [20]. When transcripts did not align to any of the databases, EST Scan (http://myhits.isb-sib.ch/cgi-bin/estscan) was conducted to decide its sequence direction.

Nucleic acid sequences from strawberry (*Fragaria X ananassa*), apple (*Malus x domestica*) and cherry (*Pruns. avium*) were aligned by blast of Uni-prot database used for *Rose roxburghii*. This was done in order to compare the four species based on exactly the same search parameters and database type.

### Identification and conserved motif analysis of *Rosa roxbunghii* superfamily

All genes related to MYBs were selected and clustered based on Nr annotation and made descriptions. The expression level of all allergen MYB genes in various tissues were showed. Conserved motifs shared by MYB proteins, significantly expressed in mature fruit using a threshold value of absolute log2 FC (fold change) ≥1 with FDR (False discovery rate) ≤ 0.01, were analyzed using Multiple Em for Motif Elicitation (MEME V 5.0.1 http://meme.nbcr.net/meme/cgi-bin/meme.cgi) online tool through uploading the amino acid sequences of the MYB superfamily members. The following parameter settings were used: 1R-MYB proteins, having one R; R2R3-MYB proteins, having two Rs; R1R2R3-MYB proteins, having three Rs; 4R-MYB proteins, having four Rs. Others belong to atypical MYB families were determined [21].

### Quantitative real-time PCR (qRT-PCR) assays

Twelve cDNAs encoding MYB transcriptional factor, all of which have potential roles in regulation plant development, were selected for qRT-PCR validation. Primers were designed (S1 Table) with primer premier 6. Total RNAs were isolated from leaf, stem, flower, young fruit (YF) and mature fruit (MF) using the Trizol, followed by purification with an RNA purification kit (Takara, Japan). Real tine RT-PCR was performed on a Roche LightCycler480 machine using SYBR green I, with *β-actin* as an endogenous control. Amplification was performed for 95°C for 2 min, 40 cycles with 95°C for 15 s, annealed at 58°C for 30 s, and 72°C for 30 s. The expression levels relative to the control were estimated by calculating ΔΔCt and subsequently analyzed using 2^-ΔΔCt^ method.

### Microsatellite detection

The program MISA (http://pgrc.ipksgatersleben.de/misa/) was used to detect microsatellite repeat motifs for each unigene in order to understand distributions of microsatellites (also known as SSRs) and to develop new markers in the transcriptome of *Rose roxburghii*. The number of core repeat motifs in mononucleotide, di-nucleotides, trinucleotide tetra-nucleotide, penta-nucleotide and hexa-nucleotides was counted.

## Results

### Illumina sequencing and sequence assembly

To identify more genes, RNA-seq of different tissues, leaf, stem, flower, young fruit and mature fruit, was performed with an Illumina HiSeq 2500 (Fig 1A–F). After trimming and quality filtration of the raw data, each tissue was represented by an average of 31.38 million reads. A total of 470.66 million reads were obtained for all five tissues with three repetitions. Total reads per biological condition are showed in S2 Table. For each sample, 86.21% of the reads could be mapped uniquely. A total of 212,534 transcripts were obtained by assembling clean reads using the Trinity program, with an average GC content of 42.08%, an average length of 1,077 bp and an N50 length of 2,085 bp. Transcripts were further clustered and assembled into 63,727 unigenes. The unigenes were all longer than 200 bp, with average and N50 lengths of 995 bp and 1,895 bp, respectively (Table 1). Of the 63,727 unigenes, 78.03% (49,727) were longer than 600 bp and 56.07% (35,732) were longer than 1 kb (Fig 2). In addition, most unigenes (60,901) had a length of less than 5,200 bp (95.57%) (S3 Table and Table 1).

**Fig 1.**
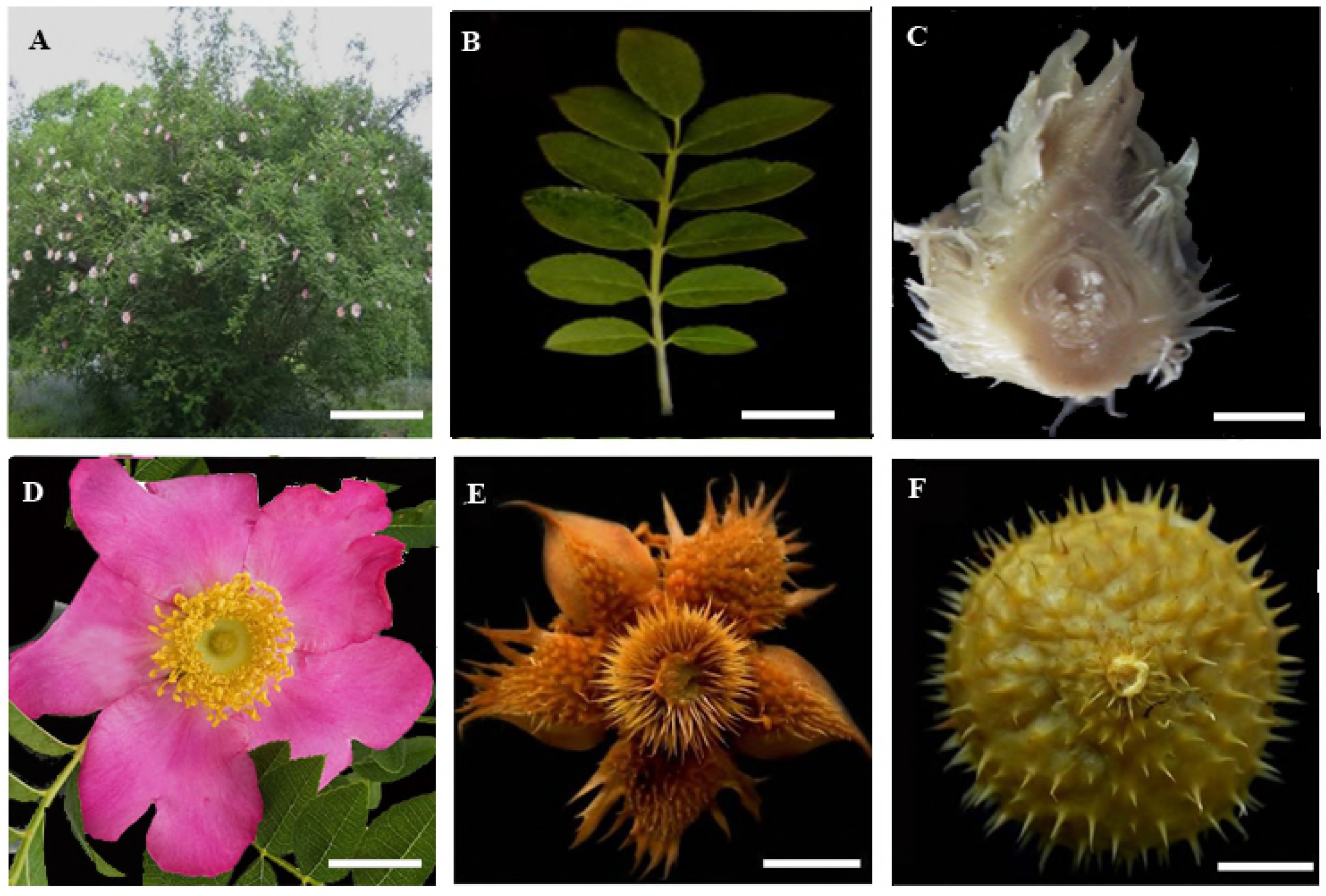
*Rosa roxbunghii* plant and different tissues collected in this study. (A) The whole plant, Bar = 50 cm; (B) Leaf and stem, Bar = 0.5 cm; (C) Flower bud, Bar = 1mm; (D) Flower, Bar =1 cm; (E) Young fruit (50 days after flowering, DAF), Bar = 1 cm; (F) Mature fruit (120 DAF), Bar =1 cm.

**Fig 2.**
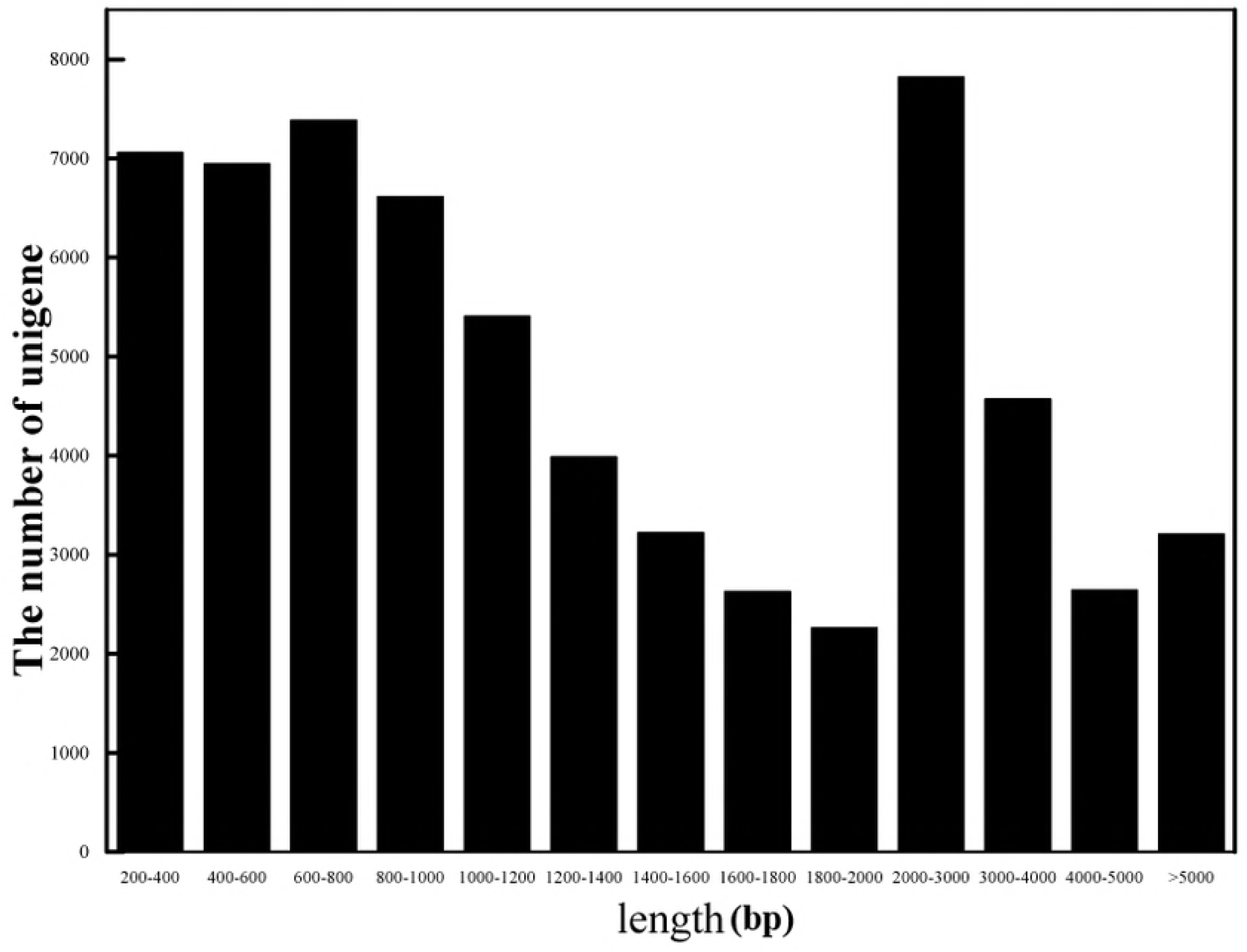
Length distribution of unigenes assembly for *Rosa roxbunghii* Tratt.

**Table 1.**
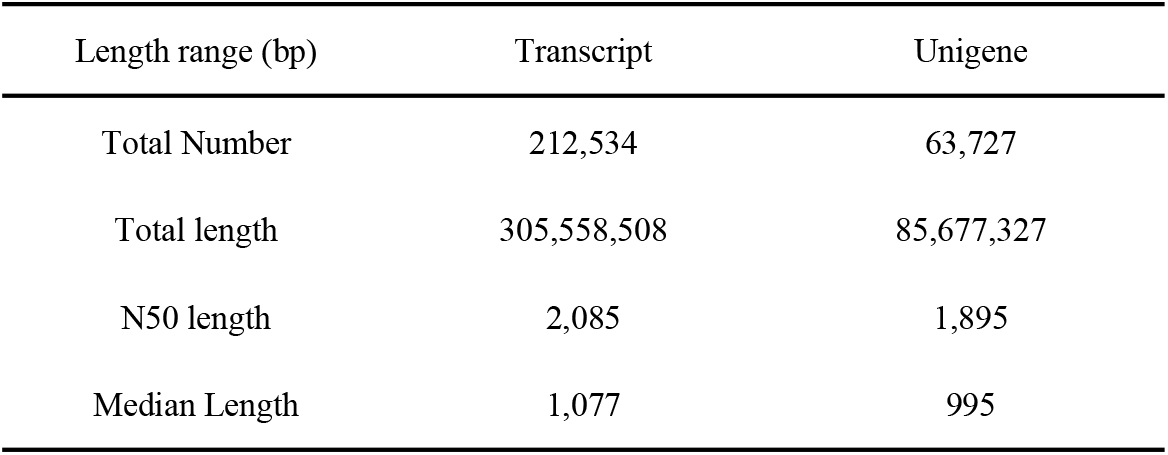
Summary of the trinity assembly for *Rosa roxbunghii* Tratt.

| Length range (bp) | Transcript | Unigene |
| --- | --- | --- |
| Total Number | 212,534 | 63,727 |
| Total length | 305,558,508 | 85,677,327 |
| N50 length | 2,085 | 1,895 |
| Median Length | 1,077 | 995 |

### Functional annotation of unigenes

To obtain sequence information, GO, KEGG, Nr, Swiss-prot, and Eukaryotic Orthologous Groups (KOG) databases were used to compare 59,358 (93.14%) of the 63,727 unigenes using a BLASTX-based algorithm (S4 Table). Swiss-prot and Nr contained the most homologous unigenes, at 55,118 and 55,151, respectively. In total, 3,284 unigenes were annotated to all database (Fig 3A). Comparative approaches effectively found differences and similarities among different species. Nucleic acid sequences from *Rose* species, including strawberry, apple, and cherry, were aligned using a BLAST algorithm-based search of the Uniprot database. We found that strawberry was the closest model species, followed by cherry and then apple, with the numbers of unigenes similar to those of *R. roxburghii* being 30,577 (47.98%), 20,105 (31.55%), and 17,677 (27.74%), respectively (Fig 3B).

**Fig 3.**
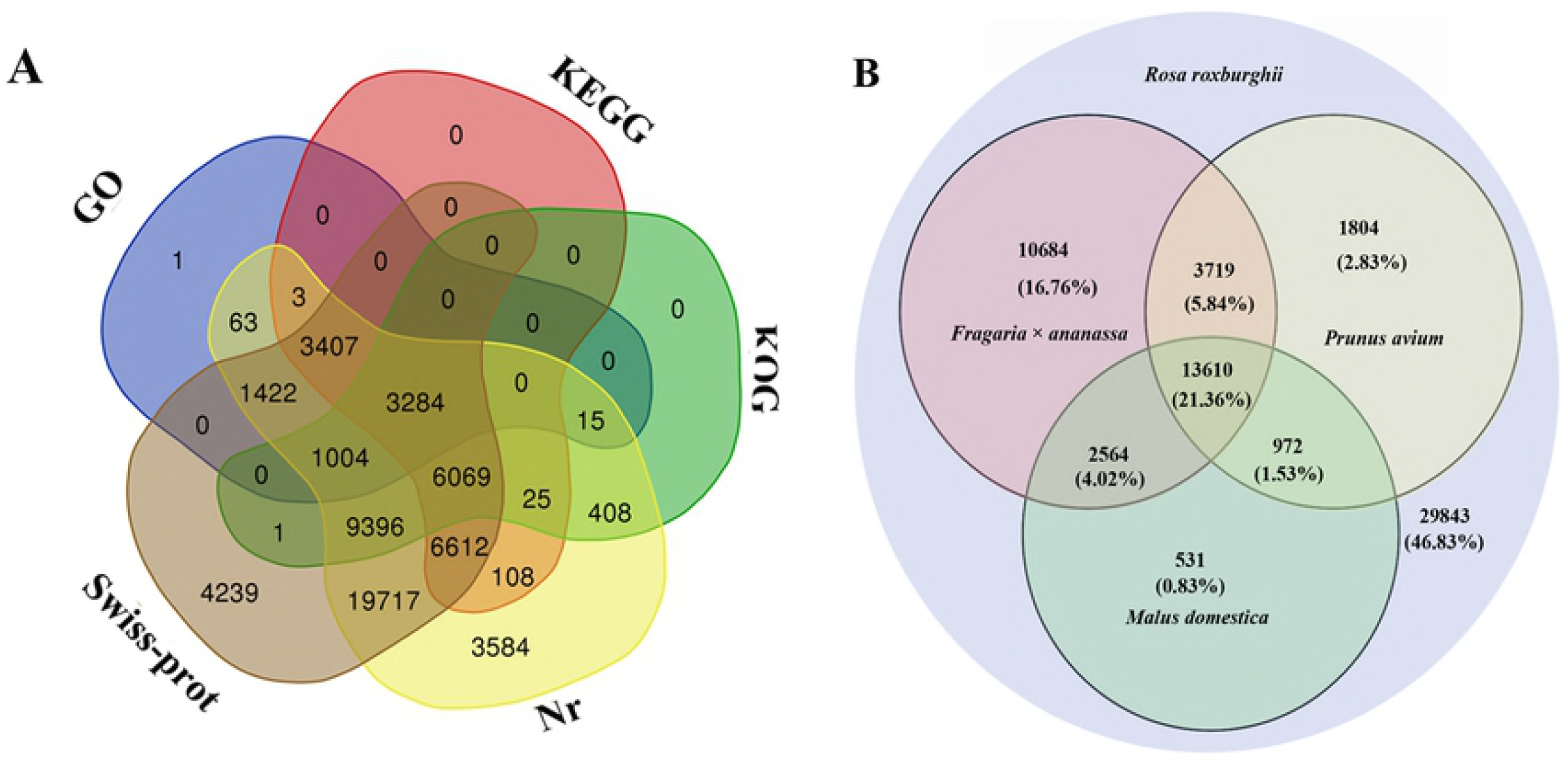
Overview of functional annotation for unigenes of *Rosa roxbunghii* (A) Venn diagram of the number of unigenes annotated by BLASTx against five different databases. The number in the circles represented the number of unigenes annotated by single or multiple databases; (B) Homology to other model Rosa species, including strawberry (*Fragaria X ananassa*), apple (*Malus domestica*) and cherry (*Pruns avium*).

There are three GO categories: biological process, cellular component, and molecular function (S5 Table). Category “biological process” consisted of 20 functional groups, with the major groups, metabolic process (56.56%) and cellular process (54.02%), having the same and higher numbers of annotations, respectively. For the category “cellular part”, 16 groups were predicted. Cell, cell part, and organelle were the three major groups. For “molecular function”, binding (49.02%) and catalytic activity (46.01%) were the dominant groups, followed by structural molecule activity (14.89%) (Fig 4).

**Fig 4.**
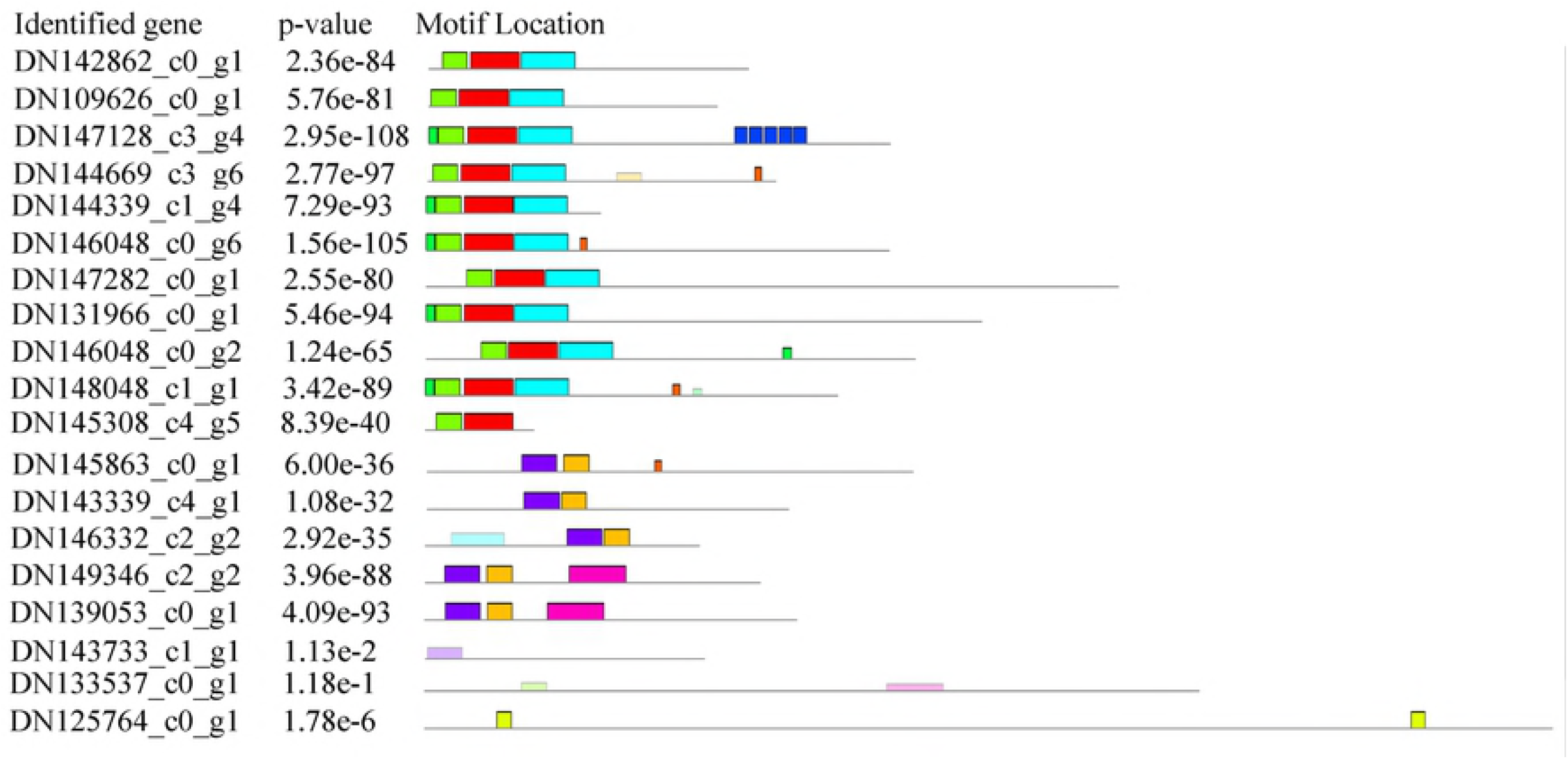
(A) Histogram of Gene Ontology (GO) classification of assembled unigenes. Blue indicated biological process, green indicated cellular process, and red represented molecular function. (B) Pathway assignment based on KEGG analysis. The bottom x-axis indicated the number of unigenes in a specific category. The y-axis indicates the clustered function groups, the right y-axis indicates the specific category of genes in the main category.

A total of 20,202 unigenes were identified using the KOG database (S6. Fig) and annotated to 25 functional categories. General function prediction (46.37%) was the largest group, followed by signal transduction mechanisms (24.26%), posttranslational modification, protein turnover, and chaperones (23.49%), and translation, ribosomal structure, and biogenesis (19.80%). The numbers of unigenes assigned to transcription (11.81%), carbohydrate transport and metabolism (11.70%), energy production and conversion (11.67%), and intracellular trafficking, secretion, and vesicular transport (11.29%) were almost the same. In addition, lipid transport and metabolism represented 10.29%. However, there were still 1,939 unigenes with unknown functions. KOG classifications revealed potential biological functions and provided an insight into the chemical reactions involved in the molecular processes in *R. roxbunghii*.

A total of 19,507 annotated unigenes were assayed to determine the biological pathways represented in *R. roxbunghii*. Briefly, these unigenes matched 140 KEGG pathways, as summarized in S7 Table. Translation in genetic information processing (3,005) was the dominant pathway, followed by carbohydrate metabolism (1,988), folding, sorting and degradation (1,375), energy metabolism (1,285), transport and catabolism (1,184), amino acid metabolism (1,161), and lipid metabolism (901) (Fig 4). KEGG pathways can provide new insights into *R. roxbunghii* biology and contribute to the prediction of the higher-level complexity of cellular processes and organismal behavior based on this information.

### Genes involved with MYB transcriptional factors in five different tissues

Among the functional database annotations, 163 MYB proteins were detected. Descriptions of the MYBs are listed in S8. Fig and S9 Table. MYBs in *R. roxburghii* were similar to the 61 members in *Fragaria vesca.* MYBs regulate secondary metabolism, gene expression and are involved in environmental stress responses. The expression levels of 159 putative *MYB* genes in five tissues are shown clustered. Various MYBs were expressed in different tissues. The black boxes represented MYBs that were significantly expressed in mature fruit.

To investigate features of homologous domains and each repeat MYB domain that was significantly expressed in mature fruit compared with the other four tissues, an online MEME was used to search for conserved motifs shared by these proteins by uploading the amino acid sequences. In total, 18 candidate *MYB* genes were analyzed using the transcriptome. Among them, 10 R2R3 *MYB,* and 1 R1 *MYB* were identified, while the remaining 7 *MYB* genes belonged to atypical MYB families (Fig.5). Sequences of the different conserved motifs are shown in S10. Fig.

**Fig 5.**
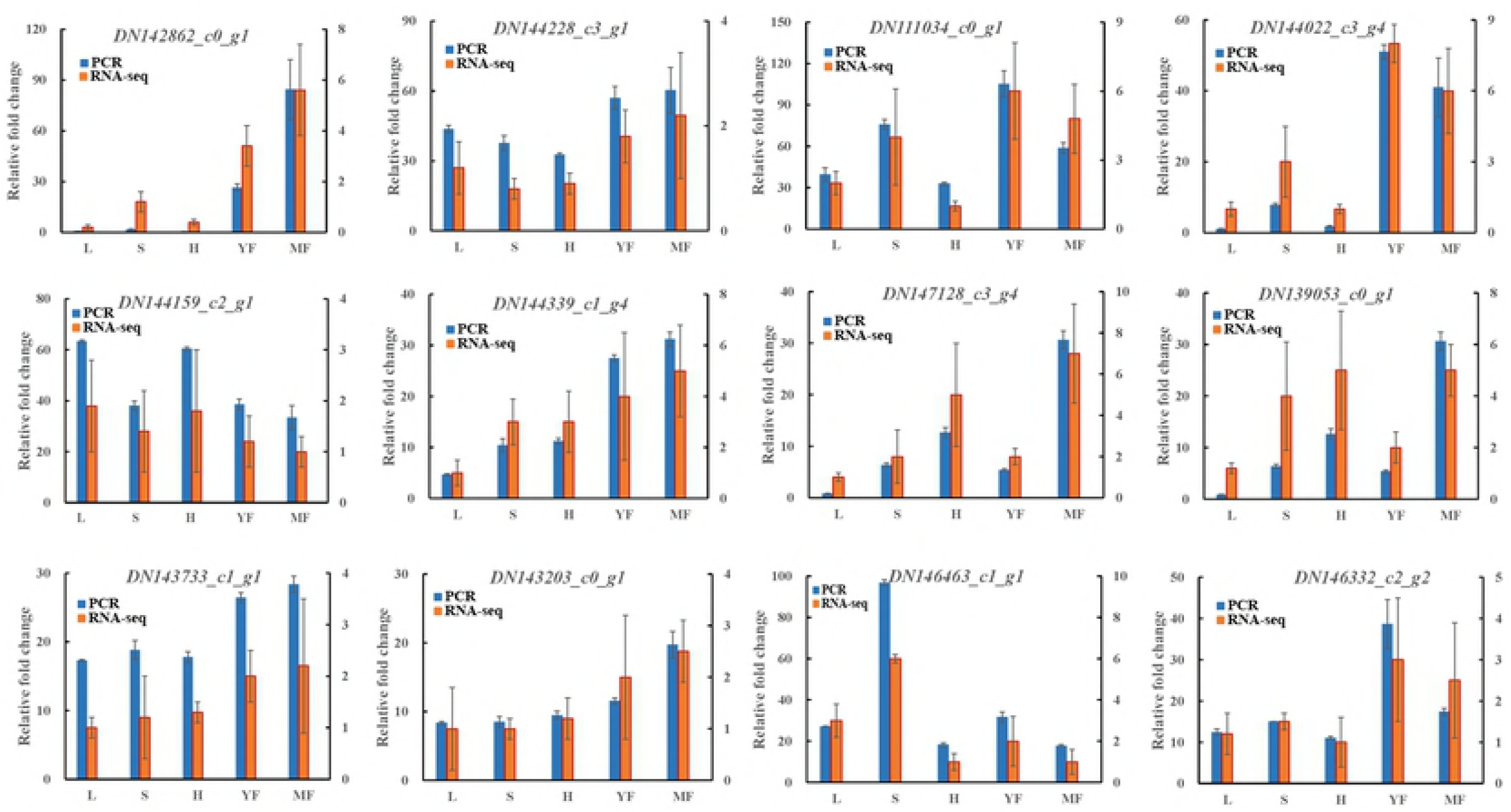
Motif distribution of MYB proteins which significantly expressed in mature fruits. Various colors represented different conserved motifs.

### Verification of RNA-seq results by qRT-PCR

Real-time RT-PCR was conducted to validate the identification of *MYB* genes by RNA-seq in five different tissues. With β-actin as the internal control, 12 genes related to MYB transcription factors were randomly selected. Validation results showed that the change trends of the 12 genes were nearly consistent with the gene expression patterns identified by RNA-seq (Fig 6). These conclusions highlight the fidelity and accuracy of the RNA-seq analysis in the present study.

**Fig 6.**
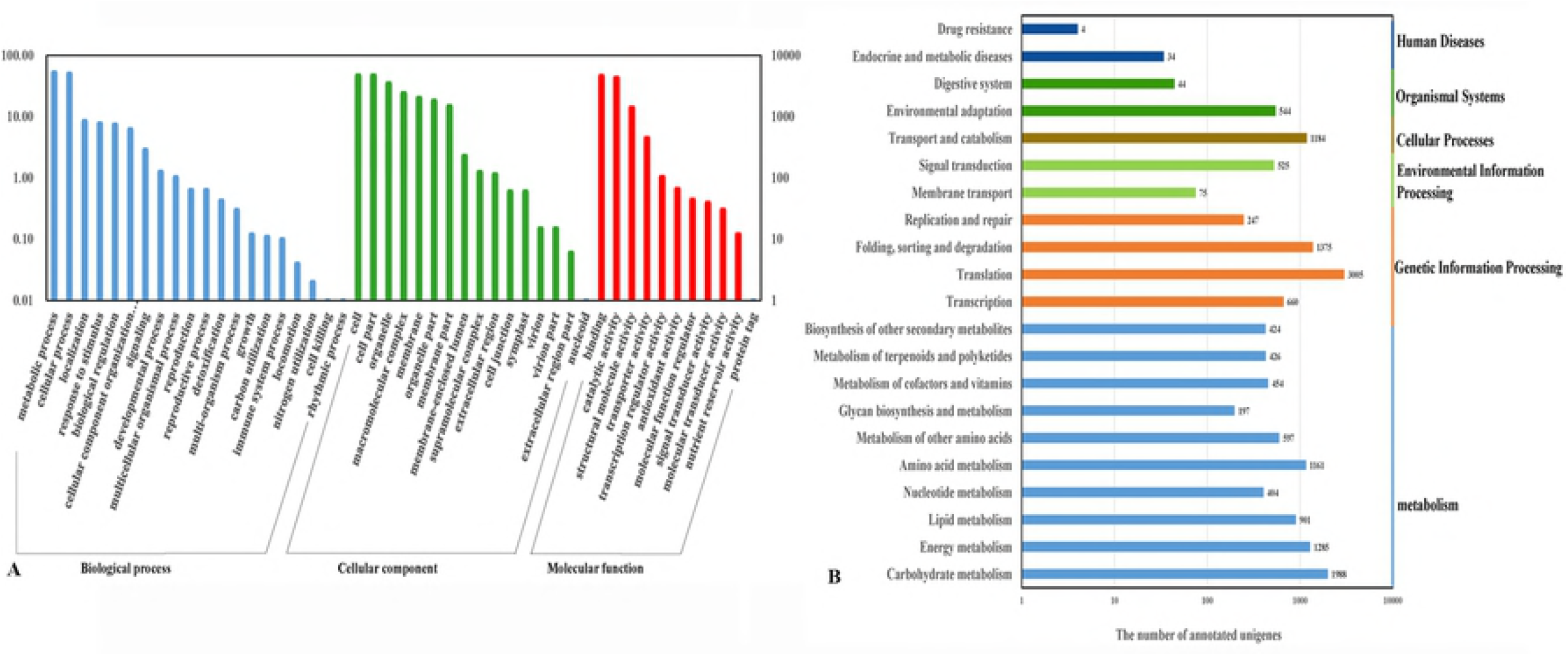
Comparison of the relative expression levels of twelve randomly selected MYB genes by RNA-Seq and RT-PCR.

### Microsatellite analysis and SSR distribution

Using MISA software, 63,727 unigenes with a total length of 109,644,660 bp were screened for microsatellite determination. A total of 37,545 potential EST–SSRs were identified. The average distribution of the SSRs was calculated to be 1:2,920 bp (37,545/109,644,660), and the average frequency of an EST–SSR was 0.59 (37,545/63,727). In total, 20,321 unigenes contained one kind of SSR (54.12%), and 8,757 contained more than one SSR (23.32%; S11 Table). In total, 5,275 (14.05%) SSRs were present in a compound formation. Transcriptome types of SSRs, from single nucleotide to hexa-nucleotide, were abundant.

Among the identified 37,545 SSRs, repeats having mononucleotide motifs were the most abundant (19,589, 52.17%), followed by di-nucleotides (11.504, 30.64%) (Table 2). The most abundant motif was A or T (18,855, 50.22%), followed by AG or CT (8,055, 21.45%) (Table 2). Among SSRs with tri-, tetra-, and penta-nucleotides, the most abundant types were AAG/CTT (2,189, 5.83%), AAAT/ATTT (141, 0.38%), and AAAAT/ATTTT (19, 0.05%), respectively. The hexa-motifs AAGGAG/CCTTCT and ACCTCC/AGGTGG, were the most abundant types and were equally present (7, 0.02%) (Table 2). The repeat positions of the SSR types were analyzed and ranged from 5 to 121. Most SSR types were repeated more than 15 times, at 19.01% (7,136), while those repeated 10 times were 17.10% (6,422) (Table 3). Except mononucleotides, the repeat numbers for most SSRs ranged from 5 to 12 (9,612, 75.9%), with only small percent being repeated more than 15 times (1,177, 9.3%).

**Table 2.**
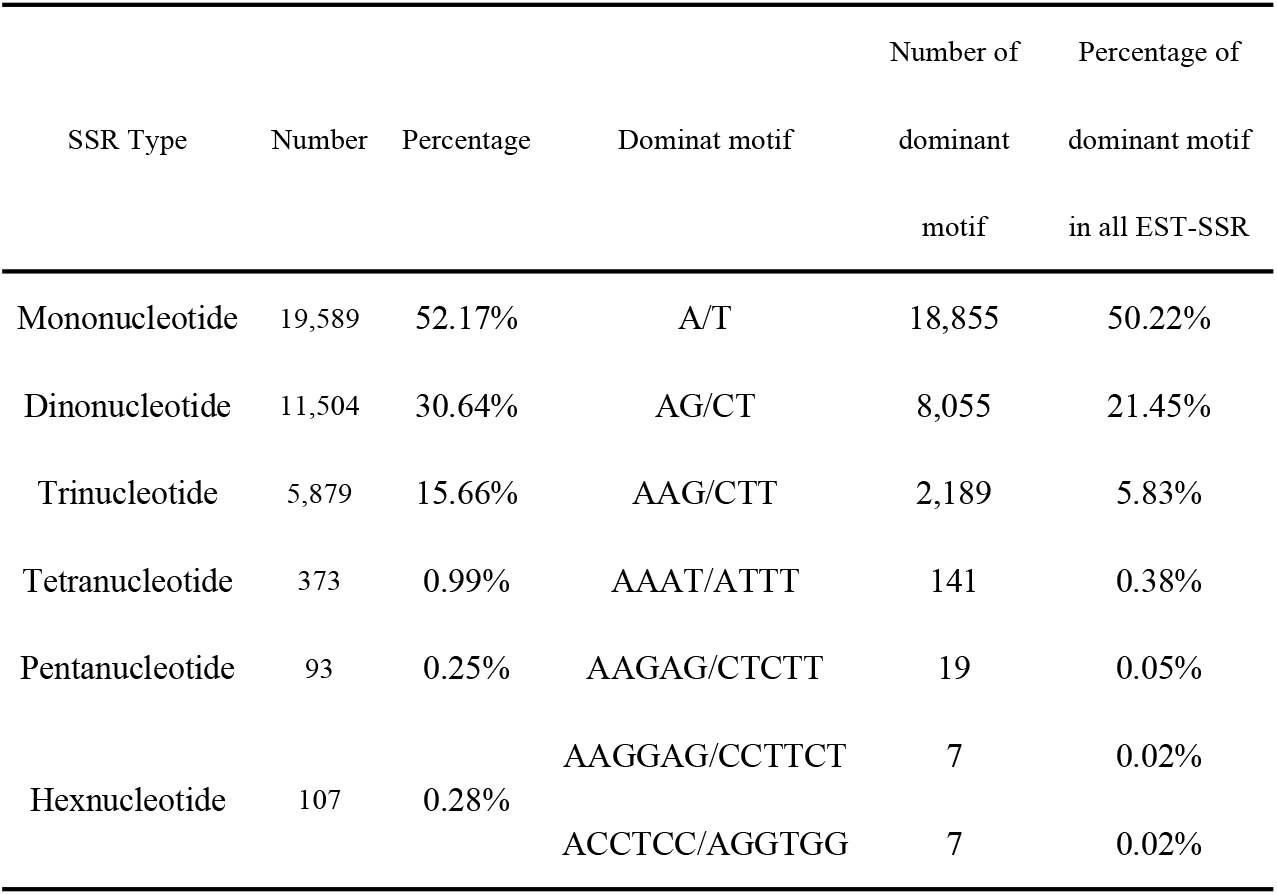
Summary of EST-SSRs identified from the transcriptome of *Rosa roxbunghii* Tratt.

**Table 3.**
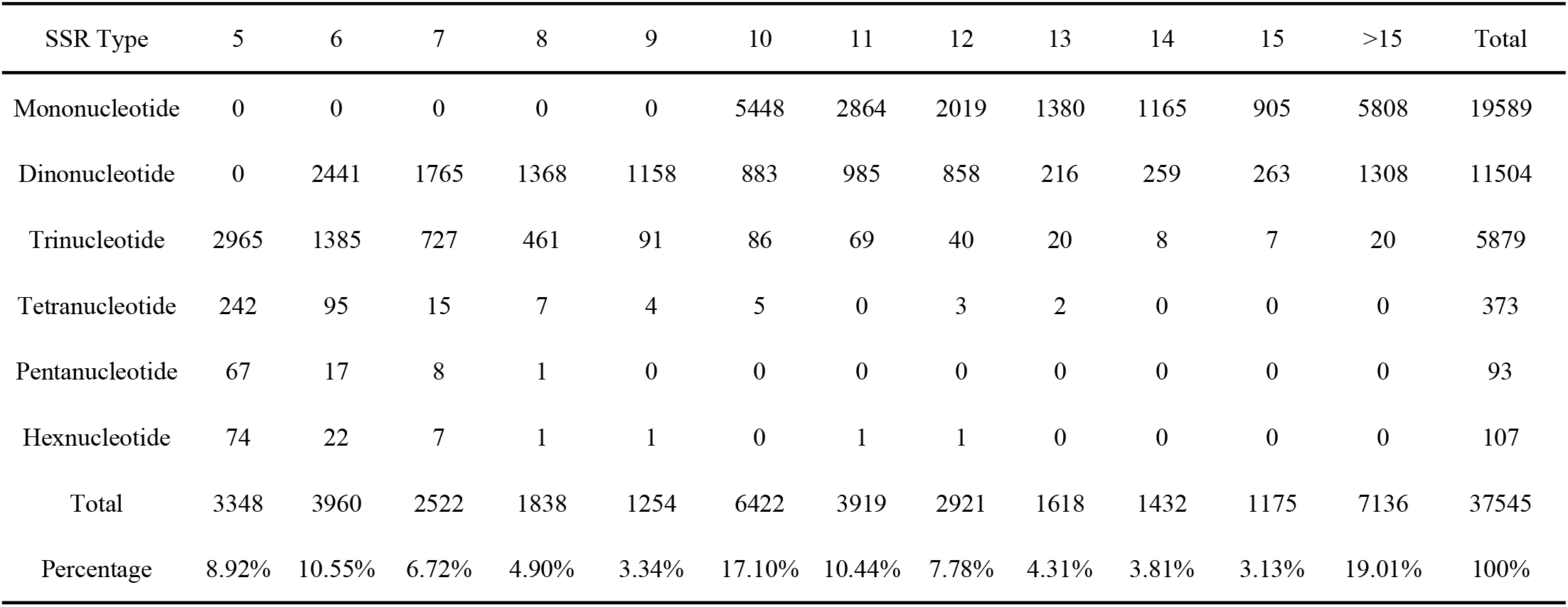
Summary of different repeat times for SSRs isolated from the transcriptome of *Rosa roxbunghii* Tratt.

## Discussion

Illumina mRNA sequencing technology is considered as an effective approach to assess transcriptional expression in different tissues. However, studies of the *R. roxburghii* transcriptome are limited. A previous study resulted in 53 million reads using fruit at three developmental stages [15]. Genomic information for *R. roxburghii* using next-generation sequencing technologies has also been obtained. In total, 30.29 Gb of sequence data was identified by HiSeq 2500 sequencing [5], covering ~60.60% of the *R. roxburghii* genome. However, whole-genome sequencing was not performed, and only genes related to the biosynthesis of ascorbic acid were the focus. The limited available genomic information for *R. roxburghii* has constrained genetic studies. To obtain sequences of as many expressed genes as possible, 15 mixed tissues were used for library construction and 470,657,040 clean reads were obtained, which was greater than previously reported. In addition, 9,354 unigenes were assigned to GO categories, compared with 9,301 unigenes in a previous study. Moreover, 19,507 unigenes were assigned to 140 KEGG pathways compared with the previous 7,480 unigenes assigned to 124 pathways. The greater amount of data increased the coverage depth and accuracy, and a large number of genes involved in different metabolic pathways can now be identified [22].

Because 15 tissues were applied to construct the cDNA libraries, a large portion of the whole transcriptome was available for annotation. In fact, 93.14% of the *R roxburghii* unigenes were annotated through BLAST algorithm-based searches against five databases. A similarity analysis in the Nr database revealed that unigenes were more homologous to cDNA sequences of strawberry than the two Rosaceae species apple and cherry. This indicated that *R. roxburghii* has a closer evolutionary relationship with strawberry. The genomic resources of strawberry have been available on line (http://bioinformatics.towson.edu/strawberry/). Strawberry has a short life cycle (3.5 to 4 months) and a facile transformation system, while *R. roxburghii* is a deciduous horticultural shrub with a long-life cycle. However, it is still feasible to study *R. roxburghii* gene functions using strawberry as a model system, as was previously done for fruit crops in Rosaceae, including apple, peach, and cherry [23].

Based on MYB expression patterns, 18 *MYB* genes were significantly expressed in mature fruit compared with in other tissues. In total, 10 R2R3 *MYBs* were identified, and this was the dominant *MYB* type. MYBs presented similar patterns and conserved motifs, suggesting that their conserved features play similar roles in group-specific functions. As part of a myb-bHLH-WD40 complex, they are involved in plant secondary metabolism and responses to abiotic and biological stresses [24, 25]. Several R2R3 MYBs were analyzed for their possible involvement in anthocyanin biosynthesis and accumulation, and fructan exohydrolase regulation when subjected to different stress and hormone treatments [26, 27]. MYBs were implicated in sugar signaling, fruit-skin coloration, and phenylpropanoid metabolism [28–30]. Most R2R3 MYBs were implicated in fruit development and should be studied to improve fruit quality and stress resistance. The results of this study will aid functional studies of interesting *MYB* genes involved in *R. roxburghii* fruit development.

Using qRT-PCR, the accuracy of RNA-seq was validated. Both assays were limited to the transcriptional level. Although 15 tissues were used to construct cDNA libraries, the whole *R. roxburghii* transcriptome was still not completely covered because some transcripts, not expressed in these tissues, may be missed.

A total of 37,545 microsatellites with different repeat types were detected from 63,727 unigenes, indicating that each unigene, on average, contained 0.59 SSR. SSR locus density was 1:2,920 bp, compared with 1:4.00 kb, in a previous study. Various criteria and parameters for SSR detection, as well as the diversity of genomic structures and compositions can influence SSR density [31]. The differences found here might reflect genome size [32], which implied that we obtained more abundant data. Errors in sequencing and assembly mistakes that resulted in mononucleotide SSRs were relatively low [33]. Except mononucleotides, the most common SSR motif was dinucleotide repeats in the transcriptome, which was similar to previously reported results. Here, AC/GT was the most common type. In conclusion, detected EST–SSRs (37,545) were more associated with functional genes and could provide valuable genetic and genomic analyses. These EST–SSRs will be useful for genetic linkage mapping construction, comparative mapping and MAS breeding.

Taken together, a deep RNA-seq analysis was conducted on five tissues, and a total of 469.5 million reads were generated. In total, 63,727 unigenes were obtained using Trinity assembly, in which nearly 90% (59,358) were successfully annotated. The data provided comprehensive coverage of the *R. roxburghii* transcriptome and allowed us to identify the MYBs significantly expressed in mature fruit. Future studies are necessary to elucidate the MYBs involved in fruit quality and development. Finally, EST–SSRs were identified and different SSR repeat numbers isolated from the transcriptome were analyzed. This study provides a valuable resource for bioengineering and biological studies of *R. roxburghii*.

## Conclusions

In present study, a deep RNA-seq analysis was conducted on five tissues, and a total of 469.5 million reads were generated. In total, 63,727 unigenes were obtained using Trinity assembly, in which nearly 90% (59,358) were successfully annotated. The data provided comprehensive coverage of the *R. roxburghii* transcriptome and allowed us to identify the MYBs significantly expressed in mature fruit. Future studies are necessary to elucidate the MYBs involved in fruit quality and development. Finally, EST–SSRs were identified and different SSR repeat numbers isolated from the transcriptome were analyzed. This study provides a valuable resource for bioengineering and biological studies of *R. roxburghii*.

## Acknowledgements

This work was supported by grants from the National Natural Science Foundation of China (Grant No. 31660554, 31600214 and 31660046), Science Foundation of Guizhou Provinces (Qiankehe J zi (2015) 2117).

## Supporting information

**S1 Table. Primers for real-time PCR.**

**S2 Table. Summary of read mapping in leaf, flower, stem, young fruit and mature fruit with three repetitions.**

**S3 Table. Analysis of size distribution of unigene for *Rosa roxbunghii* Tratt.**

**S4 Table. The number of unigenes annotated according to the NCBI non-redundant (Nr), Swiss-Prot, KOG, Gene Ontology (GO) and KEGG database in *Rosa roxbunghii* Tratt.**

**S5 Table. The number of unigenes annotated with GO and classified with three categories.**

**S6 Fig. Histogram of Eukaryotic Ortholog Groups (KOG) classification of assembled unigenes.**

**S7 Table. The number of unigenes involved in KEGG pathways.**

**S8 Fig. Heat map diagram of the expression levels of MYB transcriptional factor in five different tissues. The black box represented MYBs which significantly expressed in mature fruit compared with other four tissues.**

**S9 Table. Analysis of genes related with MYB transcription factor in five different tissues of *Rosa roxbunghii* Tratt.**

**S10 Fig. The different color represented conserved motifs.**

**S11 Table. Summary of different type EST-SSRs identified from the transcriptome.**

